# Penicillin-binding protein redundancy in *Bacillus subtilis* enables growth during alkaline shock

**DOI:** 10.1101/2023.03.20.533529

**Authors:** Stephanie L. Mitchell, Daniel B. Kearns, Erin E. Carlson

## Abstract

Penicillin-binding proteins (PBPs) play critical roles in cell wall construction, cell shape, and bacterial replication. Bacteria maintain a diversity of PBPs, indicating that despite their apparent functional redundancy, there is differentiation across the PBP family. Seemingly redundant proteins can be important for enabling an organism to cope with environmental stressors. We sought to evaluate the consequence of environmental pH on PBP enzymatic activity in *Bacillus subtilis*. Our data show that a subset of *B. subtilis* PBPs change activity levels during alkaline shock and that one PBP isoform is rapidly modified to generate a smaller protein (i.e., PBP1a to PBP1b). Our results indicate that a subset of the PBPs are preferred for growth under alkaline conditions, while others are readily dispensable. Indeed, we found that this phenomenon could also be observed in *Streptococcus pneumoniae*, implying that it may be generalizable across additional bacterial species and further emphasizing the evolutionary benefit of maintaining many, seemingly redundant periplasmic enzymes.

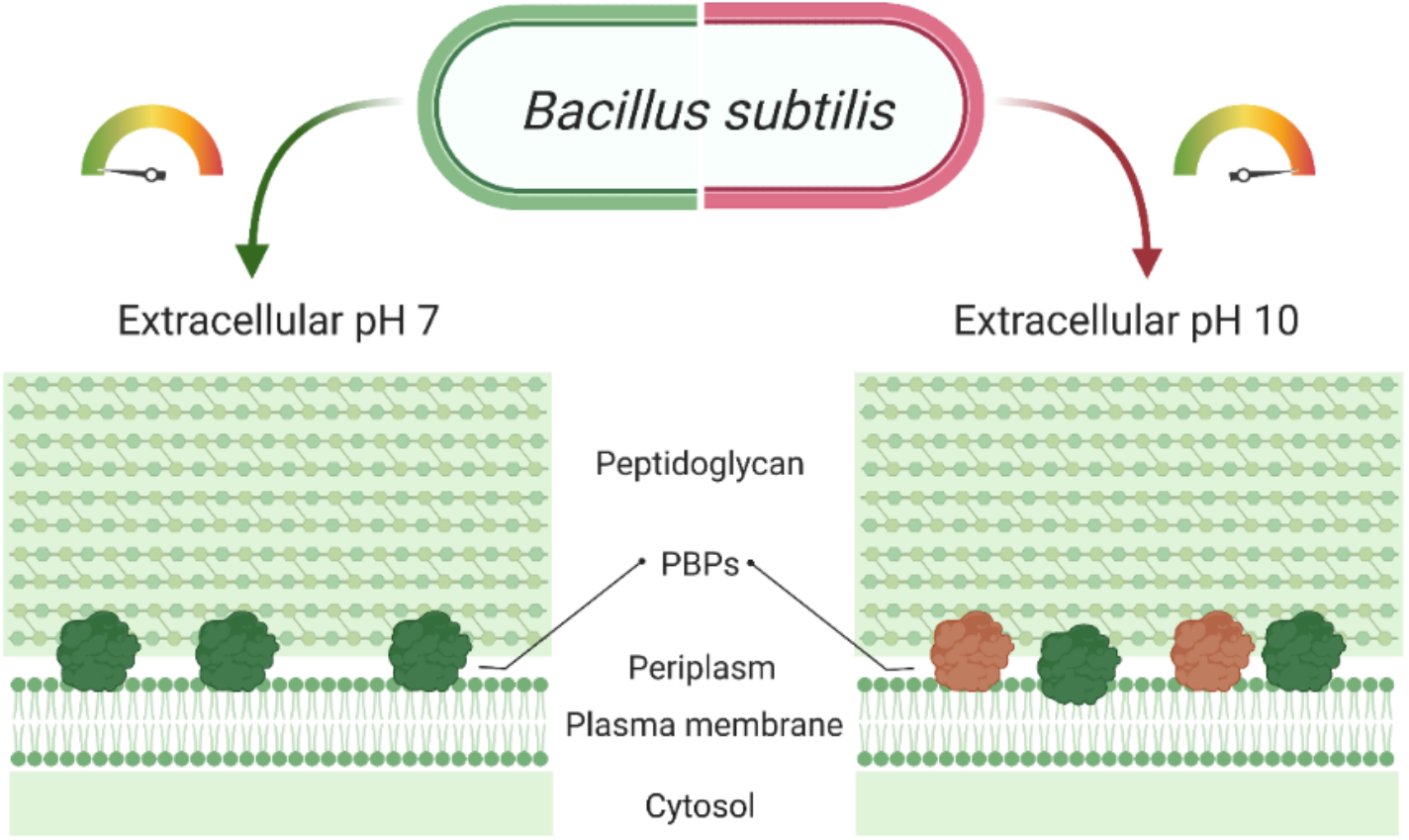

## Introduction

The evolutionary success of bacteria is largely due to their ability to sense and respond to environmental change. Bacteria, therefore, have become ubiquitous and occupy ever-changing and even extreme environmental conditions, including temperature, pH, osmolarity, nutrient, and chemical pressures. The first line of defense against changing external environments is the cellular envelope. In Gram-negative bacteria, this protective layer consists of an inner lipid membrane, a thin layer of peptidoglycan, and an outer lipid membrane. Alternatively, Gram-positive bacteria have a single, inner lipid bilayer and a much thicker peptidoglycan layer. The microbial cell envelope enables the cell to maintain cytoplasmic homeostasis and protects the internal biochemical processes from changing conditions. For example, enzyme kinetics are highly dependent on pH, so bacteria have developed several strategies for coping with fluctuations in environmental pH (*e.g*., proton pumps, other monovalent cation efflux, regulating stress response) to maintain a neutral cytoplasmic pH and enable optimal enzymatic activity (1, 2). However, both Gram-negative and Gram-positive bacteria possess proteins that are external to the protective inner membrane and are therefore subject to environmental pH fluctuations. Thus, bacteria must employ different strategies to maintain extracellular enzymatic activity, particularly those responsible for the synthesis of the essential cell envelope.

A group of key cell envelope-associated proteins are the penicillin-binding proteins (PBPs), which are responsible for the final steps of peptidoglycan biosynthesis in bacteria. Peptidoglycan, a biopolymer composed of glycan chains, is a major structural element in the bacterial envelope and provides protection from physical and chemical hazards (3). PBPs are responsible for several key reactions in peptidoglycan synthesis: polymerization of the glycan (transglycosylation) and pentapeptide crosslinking (transglycosylation). The PBPs are divided into three classes. Class A PBPs are designated as high molecular weight (HMW) proteins that contain both transglycosylase and transpeptidase domains. Class B PBPs are also HMW proteins, but only perform a transpeptidation reaction. Class C or low molecular weight (LMW) PBPs are carboxypeptidases and catalyze pentapeptide hydrolysis to prevent additional crosslinking or possess endopeptidase activity (3, 4).

PBPs are a protein group notable for their redundant activity (5). Bacteria harbor 4–16 different PBPs, indicating that the evolution and maintenance of proteins with overlapping function is advantageous. In fact, mutation of one PBP often has little physiological effect as its altered activity is masked by other isoforms (4, 6). The PBPs of numerous bacterial species have been studied extensively, with particular attention paid to the differential inhibition of PBPs by β-lactam antibiotics (6–9). However, little is understood about the role of the functional redundancy of PBPs in typical ecological environments. Extracellular enzymes tend to be more redundant than cytoplasmic proteins because of the variations in the external environment (10). It is likely that many of the PBPs are specialized for different, and largely unexplored, environmental conditions.

We sought to investigate the specialization of PBPs in *Bacillus subtilis*, a commonly studied Gram-positive, rod-shaped bacteria that has been isolated from diverse environments (11). The lifecycle of *B. subtilis* includes endospore formation, which enables the bacterium to survive extreme conditions and makes it a particularly interesting model organism for investigating bacterial replication and cell wall synthesis. *B. subtilis* harbors sixteen PBPs: four class A, six class B, and six class C. The activity of only seven of these PBPs can readily be observed during vegetative growth (6) (**Table 1**). *B. subtilis*, like most neutrophiles, is capable of replication across a wide pH range (pH 6–9) (12). Alkaliphilic *Bacillus spp*. are capable of growth in even more extreme pH (up to pH 10.8) while still maintaining a cytoplasmic pH conducive to optimal protein catalysis (1, 13, 14). Given the ability of this genus to thrive under various environmental conditions, we postulated that *B. subtilis* PBPs may be differentially regulated to enable the organism to cope with external changes. As such, we assessed the activity of *B. subtilis* PBPs *in vivo* across a range of external pH values.

**Table 1.** Summary of *B. subtilis* PBPs resolved in Bocillin-FL gel assay. The predicted molecular weight (Expasy), isoelectric point, class, localization, and function of the PBPs resolved in an activity assay (Bocillin-FL SDS-PAGE), as well as their similarity to other *B. subtilis* vegetative PBPs and *Streptococcus pneumoniae* PBPs (7). These seven PBPs are active during vegetative growth of *B. subtilis*. Although *ponA* produces two possible protein products, only the sequence and molecular weight of PBP1a is known.

| Protein | Gene | MW (kDa) | PI | Class | Location | Mutant Phenotype | Reference | Similarity to other *B. subtilis* and *S. pneumoniae* PBPs |
| --- | --- | --- | --- | --- | --- | --- | --- | --- |
| PBP1a/b | *ponA* | 99.56 | 4.94 | A | Septa | Slight length and growth rate increase | (21, 22, 57) | 2c: 64% coverage, 37% identity<br>4: 58% coverage, 33% identity<br>2d: 68% coverage, 32% identity<br>*S. pneumo* (PBP1a): 85% coverage, 38% identity<br>*S. pneumo* (PBP2a): 75% coverage, 34% identity |
| PBP2a | *pbpA* | 80.14 | 9.22 | B | Even distribution | Similar to PBPH sequence | (21, 23) | H: 94% coverage, 43% identity<br>2a has a unique 27 AA stretch that has a PI of 10 not found in PBPH<br>*S. pneumo* (PBP2b): 97% coverage, 33% identity |
| PBPH | *pbpH* | 79.35 | 8.93 | B | Even distribution | Similar to wild type, double mutant with *pbpA* is not viable | (20) | 2a: 91% coverage, 44% identity |
| PBP2b | *pbpB* | 79.32 | 8.9 | B | Septa | Essential | (5) | spoVD: 83% coverage, 34% identity<br>3: 70% coverage, 26% identity<br>*S. pneumo* (PBP2x): 96% coverage, 32% identity |
| PBP3 | *pbpC* | 74.41 | 6.24 | B | Periphery | Similar to wild type | (5, 22) | spoVD: 70% coverage, 27% identity<br>2b: 62% coverage, 27% identity |
| PBP4 | *pbpD* | 70.66 | 9.12 | A | Septa and periphery | Reduced growth rate, morphological changes | (5, 21) | 2c: 90% coverage, 34% identity<br>1: 93% coverage, 32% identity<br>2d: 91% coverage, 30% identity<br>*S. pneumo* (PBP1b): 70% coverage, 28% identity |
| PBP5 | *dacA* | 48.64 | 5.77 | C | Septa and periphery | Similar to wild type | (58) | none |

We focused our work on alkaline pH as *B. subtilis* is known to grow in conditions up to pH 10 and the *Bacillus spp*. includes several species of alkaliphiles (15). The activity changes of single, purified PBPs have been reported over a range of pH values, but to our knowledge there are no reports of how pH affects PBP activity *in vivo* (16–18). We found that during alkaline shock, PBPH and PBP4 are inactivated. We also show that base treatment promotes transition between the two products of *ponA*, PBP1a and PBP1b, with higher pH values resulting in inactivation of PBP1a by transition to the smaller PBP1b protein. The changes in PBP activity did not result in notable differences in cell morphology, indicating the benefit of PBP redundancy. PBP-null mutant *B. subtilis* strains revealed the role of alkaline-active PBPs (2a, 2b, 3, and 5) in maintaining bacterial replication in alkaline media. We also found that *S. pneumonia* possesses a base-sensitive PBP, indicating that redundancy within this protein family is likely important for survival in environments with differing pH values in multiple organisms.

## Results and Discussion

### Alkaline shock reduces the activity of select PBPs

We sought to determine if alterations in external pH would affect PBP activity in *B. subtilis*, a model organism for the study of cell wall construction. To assess the activity of these enzymes, we enlisted an activity-based probe that covalently labels proteins in proportion to their activity. As their name suggests, PBPs are the binding target of β-lactam antibiotics such as penicillin. These molecules covalently tag the PBPs, which prevents enzymatic action and halts new cell wall synthesis, thereby causing cell death. All three classes of PBPs contain a catalytic serine that interacts with the terminal D-Ala-D-Ala of the peptidoglycan pentapeptide, the moiety that β-lactams mimic (**Figure 1**). Bocillin-FL, a fluorescent analog of penicillin V, has affinity for all active PBPs and therefore can be used as an activity-based probe to report on the catalytic activity of the entire protein suite (6, 19). To determine the effect of alkalinity on PBP activity, we cultured *B. subtilis* cells in LB to early exponential phase (OD_600_ 0.4–0.6), then exposed the cells to gradients of alkaline pH in PBS for 30 min. Next, cells were incubated with Bocillin-FL to label the active PBPs. Following lysis, cell membranes were isolated, and the proteome separated via polyacrylamide gel. Gels were scanned to visualize the fluorescence intensity of the bands, which indicates the degree of activity of individual PBPs.

**Figure 1.**
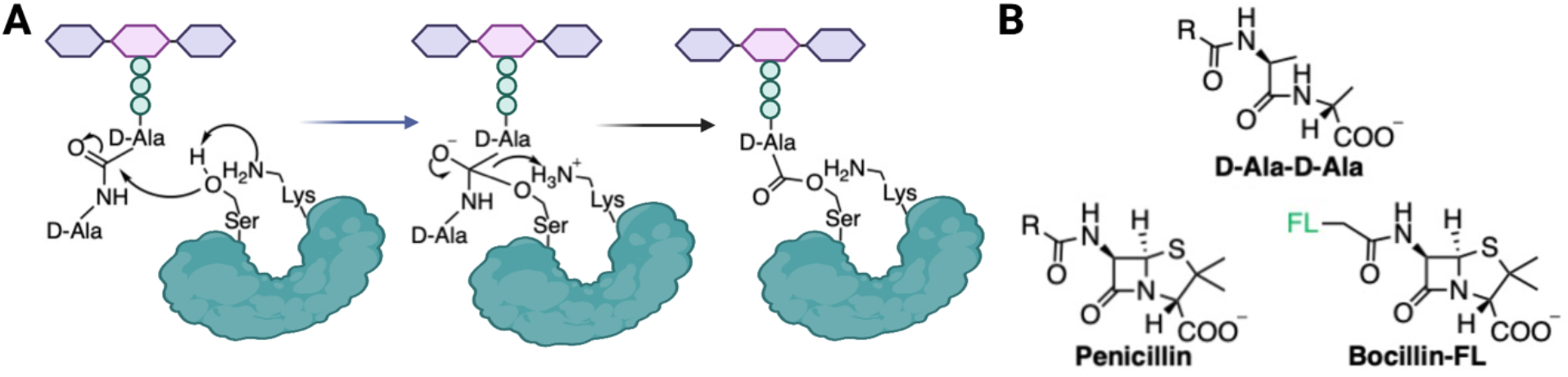
Peptidase activity and substrate mimetics of PBPs. **A**. The transpeptidase or carboxypeptidase PBP domains contain a lysine that activates serine for acylation by the D-Ala-D-Ala of the stem peptide. **B**. β-Lactam antibiotics, such as penicillin, mimic this natural substrate. A β-lactam conjugated a fluorophore (FL) can be employed to assess the activity and localization of the PBPs.

We found that with increasing pH, there was a loss of activity of PBPH, PBP4, and a shift from PBP1a to PBP1b as indicated by the disappearance of fluorescent signal due to the reduction in labeling of the activity-based probe, Bocillin-FL (**Figure 2**). PBPH (*pbpH*), which is more active in the later log phase of vegetative *B. subtilis* growth and is evenly distributed across the bacterial membrane, is fully inactivated at the modest pH of 8.5 (20). PBPH and PBP2a (*pbpA*) have the highest sequence and functional homology among the PBPs in *B. subtilis* (**Table 1**). *B. subtilis* strains lacking either of these genes do not display morphological defects, although mutants without both PBPs (*ΔpbpHΔpbpA*) are not viable. Despite sequence similarity with PBPH, PBP2a activity is not affected by increased alkalinity. This may suggest that the redundancy of PBPH and PBP2a facilitates growth in a wider pH range as PBP2a appears to be better able to function under alkaline conditions.

**Figure 1.**
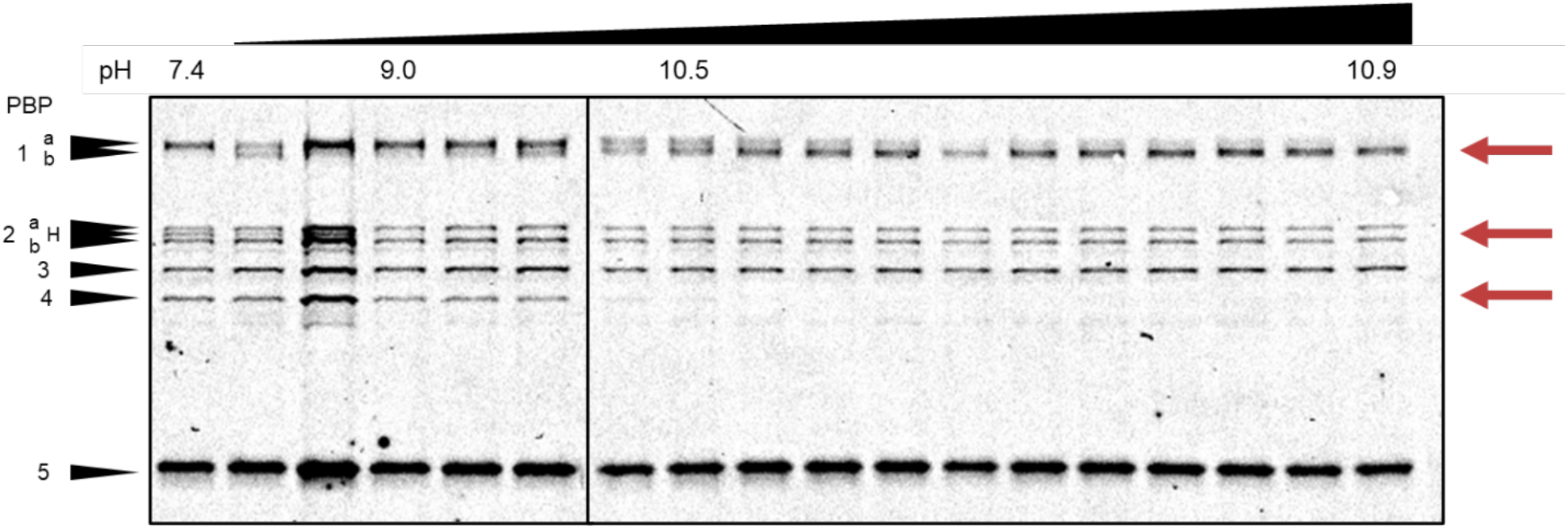
Inactivation of PBPH, PBP4, and activity shift from PBP1a to PBP1b following treatment with NaOH. Representative SDS-PAGE gel image for titration of *B. subtilis* 3610 cells over the indicated pH range. The fluorescent signal is a result of degree of labeling with Bocillin-FL, an activity-base probe. Red arrows indicate inactivation of noted PBPs.

PBP4 (*pbpD*) inactivation begins at pH 10.5 (**Figure 2**). PBP4 is active at the septa and periphery of *B. subtilis* during vegetative growth. *B. subtilis* strains lacking PBP4 (Δ*pbpD*) have a reduced growth rate and slight morphological changes (5, 21). When compared to other PBPs in *B. subtilis*, PBP4 is most homologous to other class A PBPs: vegetative PBP1 (*ponA*) and sporeforming PBPs, PBP2c (*pbpF*) and PBP2d (*pbpG*), the latter of which are not observed in this assay as it was performed during vegetative growth (**Table 1**). Therefore, it is unclear if the activity of the spore-forming PBPs are affected by an alkaline environment. However, we do also see a substantial change in PBP1 activity (see below), which aligns with the predicted similarity between PBP1 and PBP4.

The final observed change in PBP activity was a shift from PBP1a to PBP1b at pH 10.5 (**Figure 2**). PBP1a and PBP1b are two products of different molecular weights encoded by the same gene (*ponA*). The biological process that results in two functional proteins of different sizes from *ponA* is not well understood but is suggested to be due to differential *C*-terminus processing (22). *B. subtilis* strains lacking *ponA* (Δ*ponA*) result in a loss of both PBP1a and PBP1b isoforms and cause minor morphological changes such as a reduction in cell diameter, cell bending, and decreased sporulation efficiency (**Table 1**) (23). Interestingly, PBP1a activity is generally dominant at neutral pH, but during alkaline shock PBP1b is activated with almost no PBP1a activity detected. Other research has shown the role of PBPs in enabling cell wall synthesis over a range of environmental pH or the changes in PBP activity across a pH range *in vitro*, but to the best of our knowledge, this work is the first report of *in vivo* PBP activity changes due to pH perturbations (24). We also conducted a preliminary investigation of the sensitivity of *B. subtilis* PBPs to acidic conditions. We found that PBPH was also sensitive to acidic conditions, as was PBP2a (**Figure S1**). Future work will more thoroughly evaluate this parallel result.

To evaluate any differences between *in vivo* and *in vitro* PBP activity and to determine if the based-mediated PBP (de)activation events resulted from a limitation of the assay instead of being a genuine biological result, we altered the order of the steps (*i.e*., base treatment, cell lysis, Bocillin-FL labeling). *In vitro* analysis was performed on *B. subtilis* lysates, which were treated with base, washed, and then Bocillin-FL labeled. Bocillin-FL labeling of *B. subtilis* lysates does not produce the same PBP profile as labeling on the whole cell, both PBP2a and PBPH remain unlabeled in lysate control samples, so we are unable to make conclusions about the regulation of PBPH (**Figure 3a**). However, these data do show that PBP4 is inactivated around pH 10.5, indicating that the loss of PBP4 activity does not require the native environment of the cell and is more likely due to biochemical changes such as the ionization state of amino acids or variations in protein folding. We also observed a decrease in the activity of PBP1a but no activation of PBP1b, implying that PBP1b activation requires additional cellular machinery such as enzymes to generate PBP1b from PBP1a or pH-sensing proteins that regulate PBP1b activity. This was further supported by experiments performed on cells that were first base-treated, lysed, then labeled with Bocillin-FL. Again, the activity of PBP4 was lost. However, because base exposure was now performed *in vivo*, we saw transition of PBP1a activity to PBP1b labeling, indicating that intact cells are required (**Figure 3b**). Finally, we investigated the stability of Bocillin-FL-labeled proteins to base by treating Bocillin-FL-labeled proteome with alkaline conditions. Analysis by SDS-PAGE showed a dissimilar PBP (de)activation trend, presumably due to base-promoted cleavage of the Bocillin-FL-PBP acyl enzyme conjugate (**Figure 3b**). Together, these data confirm that the observed changes in PBP activity are not due to limitation or artifacts of the assay.

**Figure 3.**
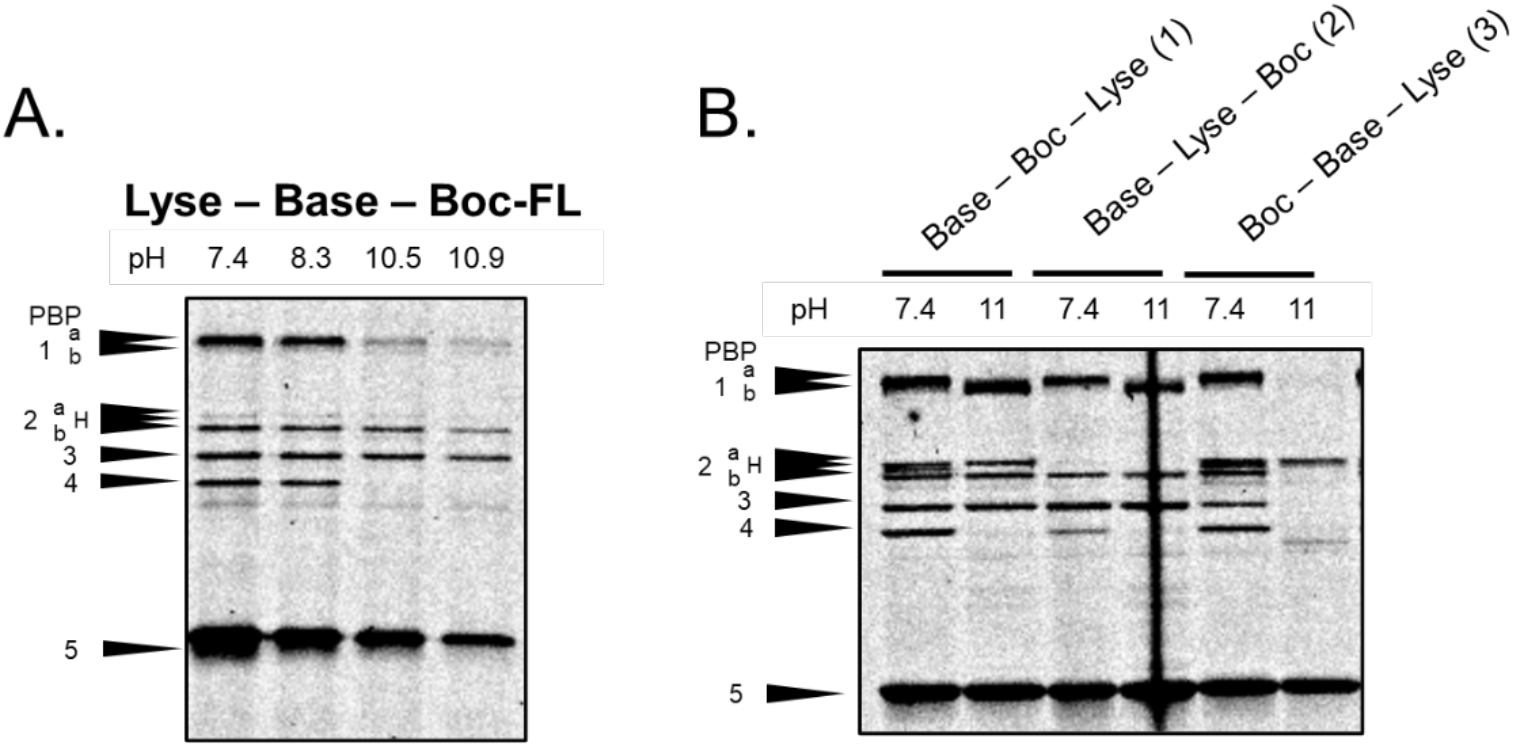
Order of alkaline shock, Bocillin-FL incubation, and cell lysis alter the activity profile of PBPs. (a) PBP profile of *B. subtilis* lysates and the effect of NaOH titration on the PBP activity. (b) Representative gel comparing the order of operations: 1) base treatment, Bocillin-FL, then lysis (standard), 2) base treatment, lysis, then Bocillin-Fl labeling, or 3) Bocillin-FL labeling, base treatment, then lysis.

To further evaluate the PBP1a to PBP1b transition, we performed a time course assay *in vivo*. These protein isoforms are encoded by the same gene (*ponA*) and are thought to result from differential posttranslational processing. PBP1a/1b are predicted to have an acidic pI, indicating their potential to be catalytically active under basic conditions (**Table 1**). Time course experiments indicate that the base-catalyzed transition from PBP1a and PBP1b requires longer than PBP4 and PBPH inactivation (10 min vs 5 min), perhaps due to regulation of the responsible posttranslational modification machinery (**Figure S2**). Like other class A PBPs, PBP1 has both transglycosylase and transpeptidase domains. Very little is understood about the difference between PBP1a and PBP1b, but Popham *et al* report that PBP1b is likely generated by truncation of the carboxy terminus region of PBP1a (22). As PBP1a and PBP1b are similar in molecular weight, PBP1b likely results from a cleavage event of less than 30 amino acids or some other more complex regulation. It has been postulated that the regulation of PBP1a and PBP1b production may be related to the presence of *prfA* (penicillin-binding protein related factor A), a gene found just upstream of *ponA* on the same operon (22). *ΔprfA* strains of *B. subtilis* display greatly reduced replication rates and abnormal cell shape (25). It is possible that alkaline shock may affect the regulation of *prfA*. However, *prfA* is also found in organisms that do not have a cell wall and additional investigation has linked *prfA* to chromosomal segregation in *B. subtilis* (25, 26).

Overall, these experiments highlight the importance of investigating protein activity and redundancy in live cells and under environmentally-relevant conditions as the apparent increase in protein processing to generate PBP1b from *ponA* would not have been identified in a more standard *in vitro* assay. Indeed, these studies provide the first known conditions that favor the activity of PBP1b over PBP1a.

### Investigation of chemical-based inhibition mechanisms

We next sought to determine the mechanism by which basic conditions reduce PBP activity *in vivo*, whether by *chemical perturbation* of the enzymatic environment (*e.g*., perturb pKa value of active site nucleophile) or *biological regulation* of PBP activity. PBPs perform three core functions (transglycosylation, transpeptidation, carboxypeptidase activity) and their active sites are uniquely tailored to each enzymatic reaction (6). The reactivity of these sites can be perturbed by the pH of the local environment, so we further investigated if changes in the catalytic activity were due to amino acid protonation state or regulatory processes. PBPH and PBP4 (class B and class A, respectively) quickly lose activity upon base incubation (5 min), so we postulated that chemical inactivation could be responsible for their decreased activity, which agrees with the inactivation of PBP4 during alkaline shock in *B. subtilis* lysates (as above; **Figure S2**). PBPs use an active site serine as a nucleophile, which is acylated by the Bocillin-FL β-lactam probe with at least one lysine required to modulate serine reactivity (27). While it is challenging to determine the pKa values of individual amino acids *in vivo*, changes in enzymatic activity across pH indicate the reactivity of the catalytic residues and suggests that the activity of purified PBPs is controlled by the protonation state of the active site lysine (**Figure 1**) (28). Literature suggests that for optimal catalysis, the proximal lysine must be in its free base form, which should be favored under basic conditions—activating the proteins. Interestingly, our experimental data reveals a more complex picture where alkaline conditions inactivate a subset of PBPs as was observed with PBP4 and PBPH. In fact, the *in vitro* experimentally-determined pH optima for the activity of many PBPs is in the basic range (pH 9–10) with lysine pK_a_ values of 8–10 (**Table S1**) (17, 18, 29, 30).

Although increased alkalinity should benefit PBP activity through lysine deprotonation, the stability of a protein is also affected by pH. The isoelectric point (pI) is the pH at which a protein has a net neutral charge and is often when proteins are least active and most unstable, which is why most proteins do not have a pI near the pH of their environment (31). For example, alkaliphilic *Bacillus* spp. are capable of growth at pH 6.8–10.8, so it follows that their PBPs and other cell wall construction enzymes are functional under extreme alkalinity and would have more acidic pI values (**Table S2**) (1, 13–15, 32–34). We expected that the pI values of *B. subtilis* PBPs may contribute to the observed pH-dependent activity and that base-sensitive PBPs would have basic pI values. However, we found no consistent trend in the relationship between predicted pI and PBP activity under alkaline conditions (**Table 1**). Both PBPH and PBP4 have predicted pI values close to the pH at which we saw reduced activity, 8.5 and 10.5, respectively. This agrees with our prediction that activity reduction under alkaline conditions is of a chemical nature. However, the pI values do not explain the activity shift in PBP1a to PBP1b or the persistent activity of other vegetative PBPs that should be inactivated during alkaline shock (PBP2a and PBP2b). PBP2a and PBPH are the most homologous PBPs and have similar predicted pI values but different base sensitivities. The most notable difference between PBP2a and PBPH is an additional 27-amino acid sequence that is unique to PBP2a. This short sequence has a predicted pI of 10, which could be partially responsible for the different predicted pI values for the proteins and serves to further stabilize PBP2a.

We would expect that the reported pH optima and predicted pI of PBPs differ, which is true for several surveyed neutrophilic organisms (*e.g., S. aureus* PBP2, *E. coli* PBP6), while some have similar pI values and pH optima (*e.g., N. gonorrhoeae* PBP4) (**Table S1**). These reported pH optima, however, were all obtained from purified proteins so they do not necessarily correlate with the *in vivo* activity of the proteins or reflect the pH of the organism’s habitat (17).

Next, we overexpressed PBPH and PBP4 from *B. subtilis* as soluble constructs in *E. coli* to investigate differences in enzyme activity that occur in a non-native environment. We did not include PBP1 since the observed transition from PBP1a to PBP1b requires protein processing. *E. coli* strains harboring plasmids for the expression of these *B. subtilis* PBPs were lysed, and the proteome exposed to base before Bocillin-FL labeling. Contrary to what was observed upon treatment of *B. subtilis* cells, we found that overexpressed PBP4 and PBPH were *activated* under basic conditions (**Figure S3**). This differential response of PBPH and PBP4 to alkaline treatment indicates a complex confluence of biological regulation, protein stability, and perturbation of active site reactivity are likely responsible for the observed PBP activity changes upon bacterial exposure to basic conditions.

### Evaluation of reagent-specific effects on base sensitivity

We next explored the generality of the observed PBP activity changes across a variety of basic reagents and buffers. Other work has shown that bicarbonate and triethylamine yield unique PBP5 activity profiles in *E. coli* (17). We found that the identity of base (e.g., lithium hydroxide, potassium hydroxide, trimethylamine) had no discernable effect on the resulting PBP activity profiles in *B. subtilis* (**Figure S4**). Several media were also tested to evaluate if buffering capacity or osmolarity stabilizes PBP activity (**Figure S5**). Alkaline shock in PBS, HEPES (2 mM), and MilliQ water all produced identical PBP activity profiles (pH >10.5). However, no PBP activity changes were observed when NaOH was added to media with higher buffering capacity (i.e., minimal media, 10 mM HEPES, 10x PBS) indicating that pH and not specific ions or ionic strength is primarily responsible for the observed deactivations. Still, salt concentration and media identity have some effect as alkaline shock in minimal media (0.25x) should alter the PBP profile as the pH is >10.5, however, we only saw decreased PBPH labeling (**Table S3**). The opposite trend was observed when alkaline shock was performed in NaCl solutions. As the concentration of NaCl increased, the PBP profile shift became more extreme, including the loss of PBP2a and PBP5 activity. Salt concentrations are known to influence base sensitivity in both neutrophiles and alkaliphiles and can compensate for some PBP-null mutants (23, 35, 36).

### Regulating pH response: Alkaline media

We next sought to determine if differential PBP activity levels under alkaline conditions had direct consequences on *B. subtilis* growth. The morphology of treated bacteria was unchanged compared to controls, which supports the benefit of PBP redundancy in maintaining the cell wall (**Figure S6a**). We analyzed the viability of the bacteria after a 30 min alkaline shock using a colony forming unit (CFU) assay. At pH 8.5 (loss of PBPH activity), bacteria are 100% viable and at pH 10.5 (loss of PBP4 activity and shift from PBP1a to PBP1b) the cells are 40% viable (**Figure S6b**).

Next, we determined if chronic exposure to basic pH would enable the organism to adapt and alter the observed PBP labeling profile. We found that *B. subtilis* was unable to survive chronic exposure to pH >9.5, similar to previously reported values (37). Chronic exposures were thus performed at pH ≤ 9.5. Overnight cultures were grown in standard LB (pH 6.65) or with the addition of NaOH. The bacteria were able to partially neutralize the media as LB is not buffered at basic pH values (e.g., pH 9.4 was neutralized to 8.0 overnight by the PBP4 mutant; **Table S4**). After overnight growth in different pH media, bacteria were subsequently exposed to a 30 min treatment with neutral media. The activity of PBPH was not recovered, while PBP1a and PBP4 activity remained inactivated (**Figure 4**; “overnight media pH” 9.4 and “alkaline shock” 7.4). Interestingly, the PBP4 activity was not lost during alkaline shock (pH 11; 30 min) if the bacteria were previously exposed to basic media (pH 9.4) overnight. This likely indicates either the presence of mechanisms to increase the stability and activity of this PBP under basic environmental conditions or that additional biological processes have been activated at higher pH to partially neutralize the periplasm and enable maintenance of PBP4.

**Figure 4.**
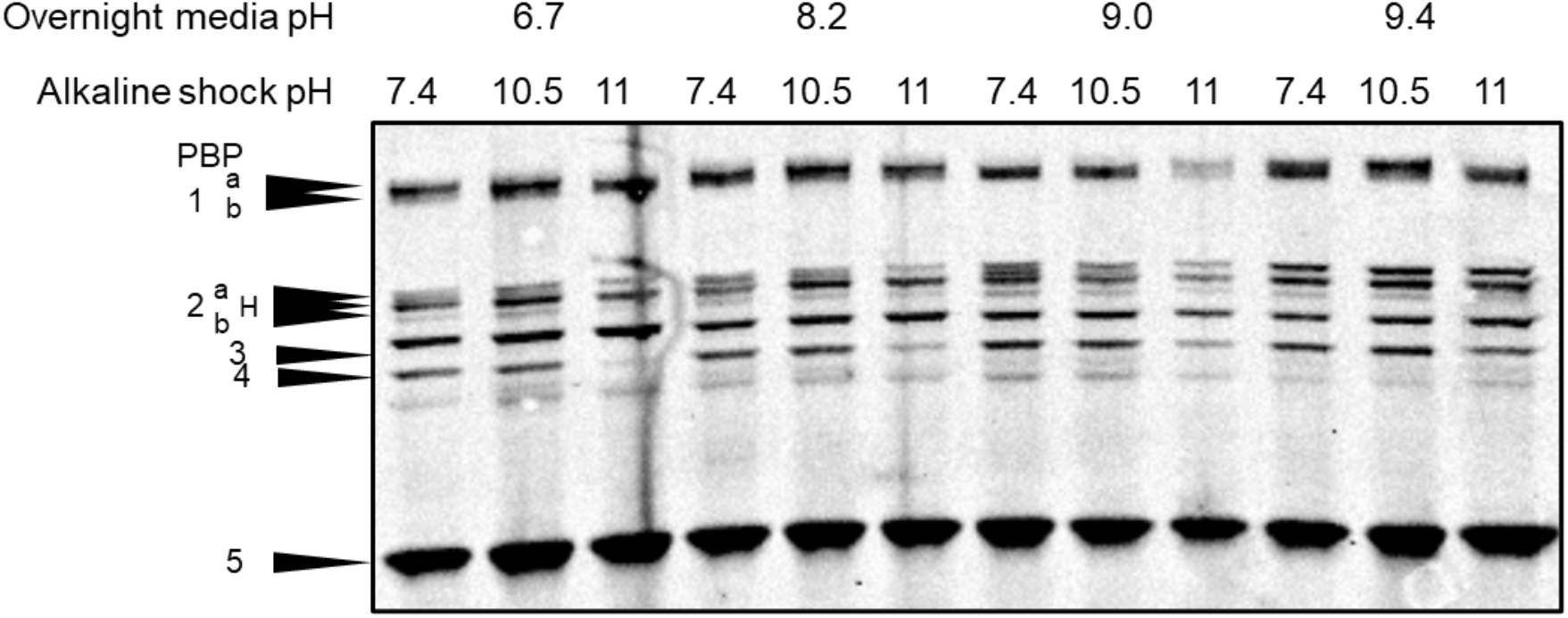
The activity of *B. subtilis* PBPs after overnight growth in basic media. Bacteria were cultured overnight in LB media that started at a pH of 6.7–9.4. *B. subtilis* were subsequently subjected to alkaline shock with at either pH 10.5 or 11.

### Alkaline sensitivity in PBP-null mutants of B. subtilis

We sought to assess the roles of specific PBPs in the ability of *B. subtilis* to survive alkaline shock. To do this, we investigated mutant organisms missing each of the vegetative PBPs, except the essential PBP2b. All mutant strains produced the same PBP profile change as the WT organism during alkaline shock (30 min): loss of PBPH activity, then PBP4 activity, and finally, transition from PBP1a to PBP1b (**Figure 5**). These data also show increased activity of PBP3 in some strains, those lacking PBPH, PBP4 or PBP5, which could indicate compensatory activity. The mutant strains also did not change morphology or Bocillin-FL localization after alkaline shock (**Figure S7**). The PBP1-null mutant, *ΔponA*, is more filamented than WT *B. subtilis* under normal growth conditions while *ΔpbpH* and *ΔpbpD* have very similar morphology to WT.

**Figure 5.**
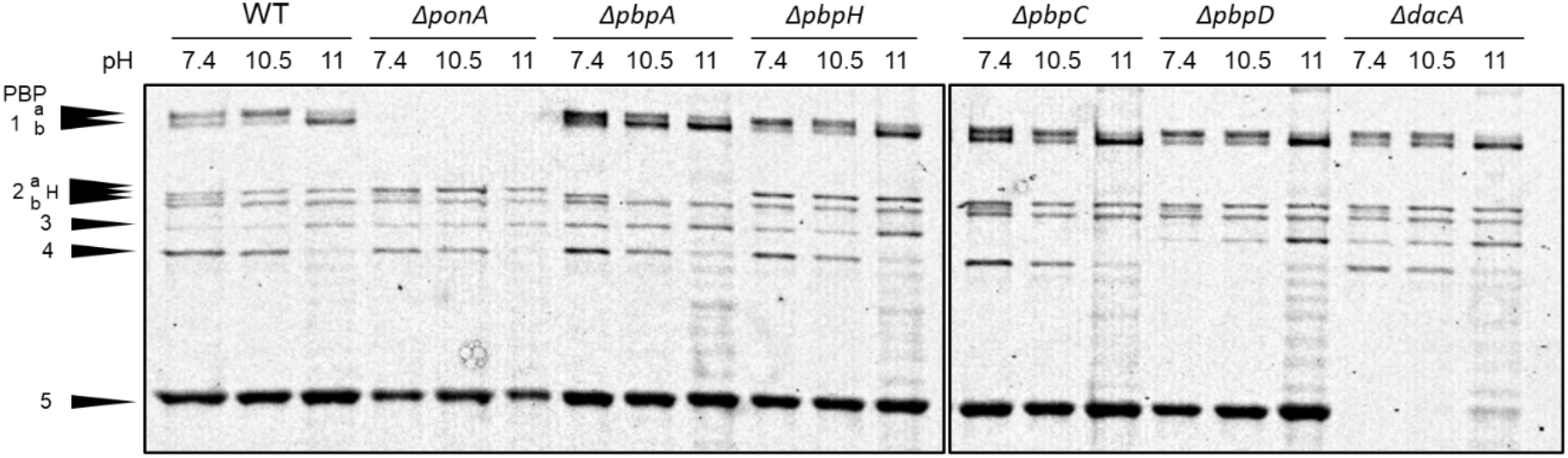
Based-mediated PBP inactivation in mutant strains. Inactivation of PBPH, PBP4, and activity shift from PBP1a to PBP1b in basic media (30 min). Representative SDS-PAGE gel image of Boc-FL labeling of *B. subtilis* 3610 cells and PBP-null mutants at pH 7.4, 10.5, or 11. WT *B. subtilis* (strain 3610), PBP1 mutant (*ΔponA*), PBP2a mutant (*ΔpbpA*), PBPH mutant (*ΔpbpH*), PBP3 mutant (*ΔpbpC*), PBP4 mutant (*ΔpbpD*), and PBP5 mutant (*ΔdacA*).

We next investigated the ability of WT and mutant strains to recover and neutralize media during long-term growth (48 hrs) in alkaline conditions to better understand the roles of individual PBPs (**Table S5**). In overnight cultures, wild-type and PBP knockout strains neutralize their media under all conditions. As expected, there was no significant difference in the ability of different mutant strains to neutralize the media as the changes in media pH are likely the result of bacterial metabolism, not the activity of PBPs (1). Growth in basic pH (pH = 9) appeared to condition both WT and mutant strains for growth at more extreme pH (pH = 9.5), as others have reported (12). However, *B. subtilis* mutants lacking alkaline-stable PBPs (PBP2a, PBP3, and PBP5) may prevent the bacteria from being able to grow at higher pH value and neutralize the media.

We next explored if the growth rate of the PBP-mutants was affected by alkaline conditions. Both wild-type and mutant strains of *B. subtilis* were cultured in LB with increasing alkalinity and growth curves were analyzed for differences in base sensitivity by comparing the amount of time required for the cultures to achieve maximal growth rate and the maximal growth rate value (**Figure 6**). Once again, LB media was not buffered to enable the bacteria to recover from the alkaline shock and neutralize the media. We calculated the maximum growth rate (peak exponential phase) by taking the first derivative of the natural log transformation of the OD_600_ measurements. Overall, we noted significant increases in the time to peak exponential phase (3–7 h) in a basic environment, which is consistent with previous reports of *B. subtilis* growth during alkaline shock (5 h growth arrest) (37). Reductions in growth rate caused by a high pH environment is likely a result of increasing cytoplasmic pH, which causes additional burden on the cells (12, 38). Because the loss of PBP activity in the mutant strains already caused growth delays, we compared the magnitude of this shift for each strain between standard growth conditions (*i.e*., neutral) and alkaline conditions. Wild-type and *ΔykuA* mutants (encodes PBPH, the most readily dispensable PBP in alkaline conditions) were the least sensitive to increasing pH values, followed by *ΔponA* and *ΔpbpD* (PBP1 and PBP4). This was expected given that the activity of these three proteins changes during acute alkaline shock, implying that they are not crucial for growth and replication under basic environments. These data also suggest that PBP1b does not provide vital functionality. The Δ*pbpC* and Δ*pbpA* strains (PBP3 and PBP2a, respectively) were more sensitive to basic pH with *ΔdacA* (PBP5) exhibiting the highest base sensitivity with the greatest increase in time to the maximum growth rate. PBP3, PBP2a, and PBP5 maintain activity during alkaline shock, so it is likely that these PBPs are required for growth in an alkaline environment. As noted previously, Δ*pbpA*Δ*pbpH* double mutants produce a lethal phenotype, so it follows that Δ*pbpA* mutants that lose PBPH activity during alkaline conditions would also have reduced cell viability. As PBP5 is the only class C PBP active during vegetative growth, it is possible that mutants without an active carboxypeptidase may be more sensitive to alkaline shock.

**Figure 6.**
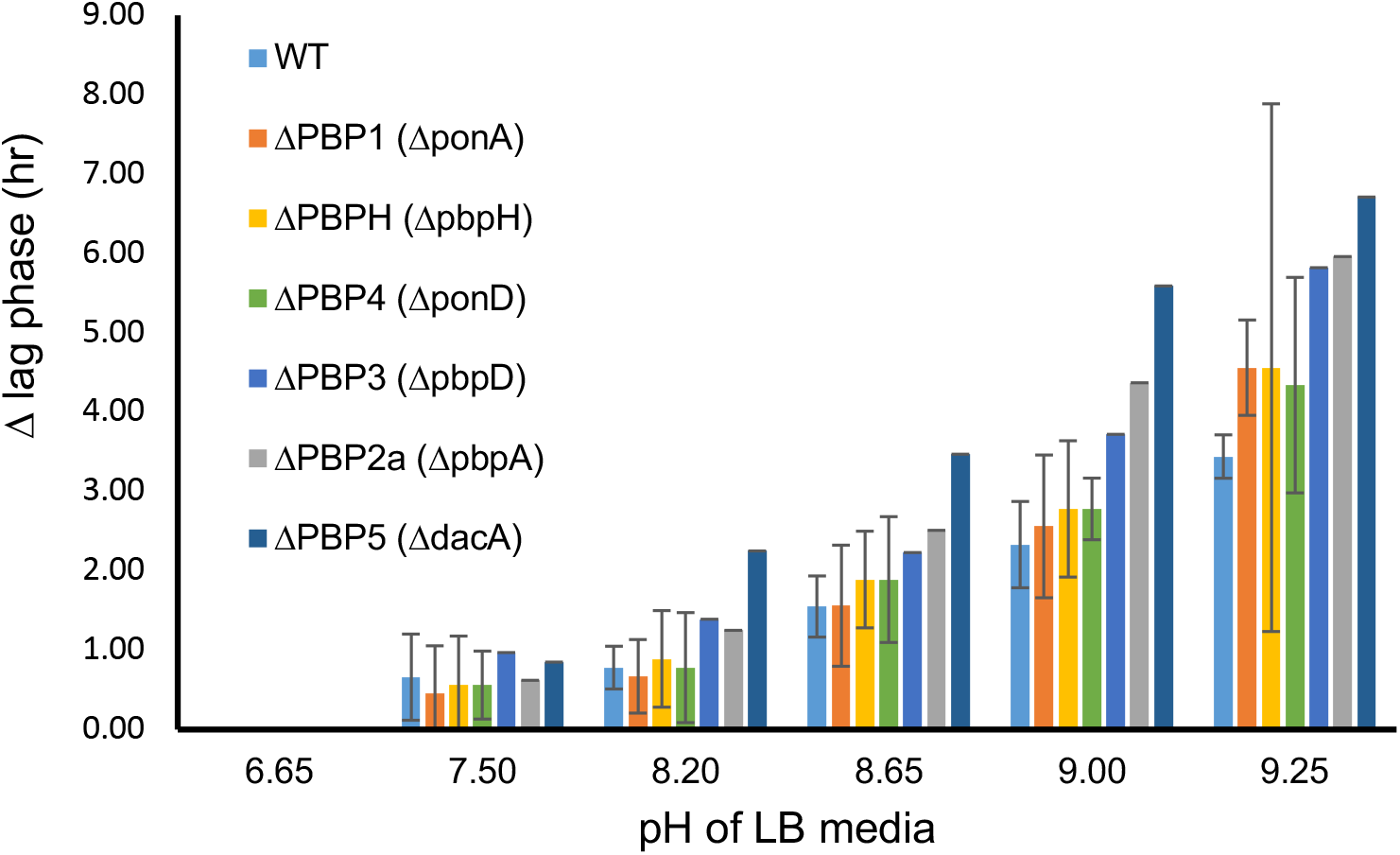
Characterization of growth curves of *B. subtilis* PBP mutants. The time for a culture to achieve maximal growth rate is considered the lag phase. The difference between the duration of the lag phase at each alkaline condition and neutral LB media is plotted for each strain.

### Additional organisms

We sought to determine if other organisms have also evolved specialized PBPs to cope with environmental pH shifts. Two additional Gram-positive bacteria were selected, *Staphylococcus aureus* and *Streptococcus pneumoniae* (39, 40). *S. aureus* and *S. pneumoniae* are both cocci-shaped bacteria that possess fewer PBPs than *B. subtilis*, suggesting less functional redundancy. Alkaline shock had no effect of the activity of the PBPs in *S. aureus* (**Figure 7a**). In *S. pneumoniae*, the activity of class A PBP, PBP1a, decreased following base treatment (**Figure 7b**). This suggests that PBP1a in *S. pneumoniae* may not be specialized for growth in alkaline conditions. PBP1a and PBP1b in *S. pneumoniae* are produced from two different genes (*pbp1a* and *pbp1b*, respectively) unlike *B. subtilis* where PBP1a and PBP1b are two functional products of the same gene (*ponA*). However, of all the *B. subtilis* PBPs, PBP1 (*ponA*) has the highest sequence similarity to PBP1a (*pbp1a*) in *S. pneumoniae* (38% protein sequence identity). The level of similarity between these two PBPs is comparable to that of *B. subtilis* PBPs that are not all affected by alkaline shock, so sequence similarity is likely not a strong predictor of basesensitivity.

**Figure 7.**
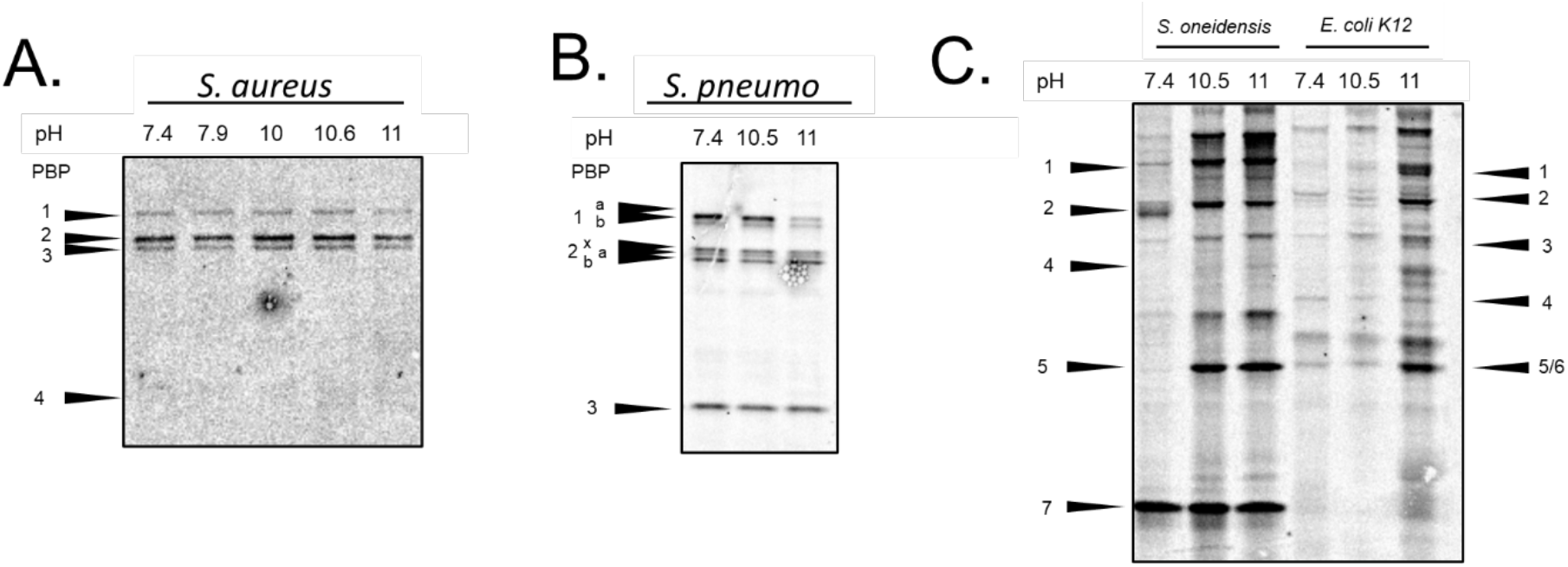
Representative gels of alkaline shock in other bacteria. (a) *S. aureus* - PBP4 does not label with Bocillin-FL, (b) *S. pneumoniae*, and (c) *S. oneidensis* and *E. coli*. The PBPs of *S. oneidensis* have not been confirmed in the literature. The presented assignments are hypothetical based on predicted molecular weight.

Gram-negative, rod-shaped bacteria *Shewanella oneidensis* and *Escherichia coli* K12 were also investigated (41). The ability of Gram-negative bacteria to maintain pH homeostasis has been well-studied, but the PBPs of Gram-negative bacteria have been historically challenging to investigate as their peptidoglycan layer is poorly accessible under the outer membrane (42). Alkaline shock increased the intensity of all bands labeled by Bocillin-FL, which suggests that the bacterial outer membrane is damaged and that there is not a specific collection of PBPs that enable growth in fluctuating environmental pH in these two microbes (**Figure 7c**).

## Conclusion

An organism’s response to environmental pH is vital for cell viability because the pH gradient across cell membranes generates energy (proton motive force) and the biochemical reactions within a cell are performed at an optimal pH (2). The activity of enzymes that maintain the bacterial cell envelope (*e.g*., PBPs) are also of interest across a range of environmental pH as they are not protected within the buffered cytoplasm. It is likely that the apparent redundancy of the PBPs is due to the importance of specialized enzymes that can withstand environmental fluctuations (24, 43). For example, in *E. coli*, only fourteen enzymes are needed to perform twelve reactions in the cytoplasmic synthesis of peptidoglycan precursors—nearly every protein performing a unique synthetic reaction. However, for peptidoglycan synthesis in the periplasmic space, there are around four enzymes available for every biosynthetic reaction (44). Mueller *et al* report the redundancy in PBPs in *E. coli* is likely needed to facilitate growth in a range of environmental pH values (24). Similarly, cytoplasmic peptidoglycan synthesis in *B. subtilis* employs numerous, discrete enzymes for unique biosynthetic reactions in contrast to the redundancy in enzymatic action in the periplasmic space (45).

The inactivation of a subset of PBPs by alkaline shock in *B. subtilis* suggests that the apparent redundancy of PBPs may enable peptidoglycan synthesis in diverse environmental conditions. These changes in activity would likely not have been resolved through predictive software or by using purified proteins. PBP1a/1b and PBP4 are both bifunctional (class A) and during alkaline stress PBP1b becomes active while PBP1a and PBP4 lose activity. Class B PBPs PBP2a, PBPH, PBP2b and PBP3 all perform transpeptidation but only PBPH is inactivated during alkaline shock. Environmental stresses such as salt concentration, osmolarity, and temperature, can be compounded by alkaline stress (1). For example, alkaliphiles require higher Na^+^ to grow at neutral pH (35, 36). Additionally, the effect of pH on bacterial replication machinery has also been linked to antibiotic resistance (2, 12, 24, 46). Finally, this work indicates that apparent functional redundancy of classes of periplasmic proteins likely indicates differential specialization for dynamic environmental conditions (10).

## Materials and Methods

### Materials and media

Bocillin-FL was purchased from ThermoFisher Scientific (Invitrogen, cat#B13233) and was stored at −80 °C. Luria-Bertani (LB) media (Lennox; 20 g/L) was autoclaved for 20 min (Sigma, powder, cat #L3022). Tryptic soy broth was prepared at 30 g/L (BD Bacto). Minimal media was prepared as a buffer (11.6 mM NaCl, 4.0 mM KCl, 1.4 mM MgCl_2_, 2.8 mM Na_2_SO_4_, 2.8 mM NH_4_Cl, 10 mM HEPES, 88.1 μM Na_2_HPO_4_, 50.5 μM CaCl_2_). Phosphate buffered saline was prepared as 137 mM NaCl, 2.7 mM KCl, 10 mM Na_2_HPO_4_, 1.8 mM KH_2_PO_4_, pH = 7.4.

### Bacterial culturing, media, exposure

*Bacillus subtilis* 3610, and *B. subtilis* PBP-null mutant strains were cultured in LB at 37 °C while shaking at 220 rpm. *E. coli* harboring the *B. subtilis* PBP plasmid constructs were also cultured in LB with ampicillin (100 μg/mL) at 37 °C while shaking at 220 rpm. *Shewanella oneidensis* MR-1 was cultured in LB at 30 °C while shaking at 250 rpm. *E. coli* K12 was also cultured in LB at 37 °C while shaking at 220 rpm. *Staphylococcus aureus* was cultured in LB at 37 °C while shaking at 220 rpm. *Streptococcus pneumoniae* was cultured in Brain Heart Infusion media at 37 °C with 5% CO_2_ without shaking. All strain identities in **Table S5**.

### pH exposures

*B. subtilis* cells grew in an overnight culture and were sub-cultured with a 1:10 dilution in fresh media until reaching an exponential culture with an OD_600_ of 0.4–0.6. A 1.0 mL aliquot was harvested by centrifugation (10,000*g* for 1 min at RT; consistent throughout protocol). Cell pellets were washed with 1 mL of PBS. Pellets were resuspended in 1 mL PBS unless otherwise indicated at indicated pH with NaOH, HCl, or other indicated solution with controls at a pH of 7.4 for 30 min at RT. Cells were once again pelleted, washed with 1 mL PBS, and resuspended in 100 μL of PBS containing 5 μM Bocillin-FL for 15 min at RT in the dark. Cells were pelleted and washed with 1 mL PBS.

### Gel-based PBP activity assay

Bocillin-FL-labeled cells were resuspended in 100 μL PBS containing 10 mg/mL lysozyme and incubated at 37 °C for 30 min. Samples were lysed on ice with a Hielscher vial tweeter UP200st (70% C, 95% A, 5% adjustment snap) for 10 min with 30 s intervals with 30 s cooling in between. The membrane proteome was collected by centrifugation at 21,000*g* for 15 min at 4 °C. The supernatant was discarded, and the membrane fraction was resuspended in 100 μL with 1% sodium dodecyl sulfate (SDS) followed by 33 μL of 4x SDS loading buffer. To denature proteins, samples were heated at 95 °C for 5 min then cooled to RT before loading 12 μL into the well of an SDS-polyacrylamide gel. Gels (10%) were prepared with acrylamide:bis-acrylamide 29:1 (Biorad) in 1.5 mm cassettes (ThermoFisher, NC2015). Proteins were resolved by gel electrophoresis at 180 V, 400 mA, 60 W for 1 h. Gels were rinsed with de-ionized water and scanned on a Typhoon 9210 gel scanner (Amersham Biosciences, Pittsburgh PA) with a 526 nm filter at 50 μm resolution. Gels were background subtracted, adjusted uniformly, and analyzed using ImageJ software (NIH).

### Microscopy

Cells were grown and exposed as described above. Cells were incubated with 3% paraformaldehyde in PBS for 25 min then washed with PBS. Fixed cells (5 μL) were pipetted onto slides (SuperFrost, FisherBrand) to air dry, then resuspended in mountant (5 μL, ProLong™ Diamond Antifade Mountant, Invitrogen) and covered with a 1.5 mm coverslip. Slides cured for 1 h and were stored in 4 °C refrigerator. Microscopy was performed with an Olympus IX73 microscope, Images were acquired with an ORCA-FLASH4.0LT+ SCMOS camera and were processed with ImageJ.

### Colony forming units

*B. subtilis* cells were grown and exposed as described above. Cells were diluted and dropped onto a plate (10 μL) in triplicate and the resultant colonies were counted after overnight growth.

### Plate reader

*B. subtilis* cells (200 μL) were cultured overnight in a Tecan Spark plate reader in LB (37 °C, shaking at 220 rpm) with 20 min timepoints.

### Marker replacement mutants

The kanamycin resistance cassette insertion-deletion constructs in *pbpA, pbpC, pbpD, pbpH*, and *dacA* were generated by isothermal “Gibson” assembly (ITA) (47). The regions upstream and downstream of each gene were amplified using DK1042 chromosomal DNA as a template and the primer pairs indicated in parentheses: *pbpA* (3534/3535; 3536/3537), *pbpC* (3538/3539; 3583/3584),*pbpD* (3598/3599; 3600/3601),*pbpH*(3579/3580; 3581/3582), and *dacA* (3542/3543; 3544/3545) (**Table S6**). Next, the kanamycin resistance gene was PCR amplified using the plasmid pDG780(48) as a template and primer pair 3250/3251. For each gene, the upstream amplicon, the downstream amplicon, and the kanamycin resistance gene amplicon were mixed in an ITA reaction, column cleaned, and transformed into *B. subtilis* strain DK1042 (49) selecting for resistance to kanamycin. Each insertion-deletion mutant was verified using PCR length polymorphism using the far upstream and downstream primers.

Isothermal assembly reaction buffer (5X) [500 mM Tris-HCL pH 7.5, 50 mM MgCl_2_, 50 mM DTT (Bio-Rad), 31.25 mM PEG-8000 (Fisher Scientific), 5.02 mM NAD (Sigma Aldrich), and 1 mM of each dNTP (New England BioLabs)] was aliquoted and stored at −80° C. An assembly master mixture was made by combining prepared 5X isothermal assembly reaction buffer (131 mM Tris-HCl, 13.1 mM MgCl_2_, 13.1 mM DTT, 8.21 mM PEG-8000, 1.32 mM NAD, and 0.26 mM each dNTP) with Phusion DNA polymerase (New England BioLabs) (0.033 units/μL), T5 exonuclease diluted 1:5 with 5X reaction buffer (New England BioLabs) (0.01 units/μL), Taq DNA ligase (New England BioLabs) (5328 units/μL), and additional dNTPs (267 μM). The master mix was aliquoted as 15 μl and stored at −80° C.

To generate the *ponA::tet* allele, the region upstream and downstream of *ponA* was PCR amplified using 3610 chromosomal DNA as a template and primer pairs 3053/3054 and 3055/3056. The tetracycline resistance gene was PCR amplified from pDG1515(48) using primer pair 2973/2974. Next, the upstream fragment was digested with BamHI-PstI, the tetracycline resistance cassette was digested with PstI-EcoRI, and the downstream fragment was digested with EcoRI-XhoI, and all three fragments were simultaneously ligated into the BamHI-XhoI sites of plasmid pUC19 to make pRC18. The plasmid pRC18 was transformed into DS2569 (49) selecting for tetracycline resistance and transduced into strain 3610 using the generalized transducing phage SPP1 (50).

### *E. coli* expression plasmids

To express the *B. subtilis* GST-PBP fusion proteins in *E. coli*, the pbpD gene was amplified from DK1042 chromosomal DNA using primer pair 7520/7521 and the pbpH gene was PCR amplified using primer pair 7522/7523 (**Table S6**). Next, the plasmid pGEX-2TK (Sigma-Aldrich) was digested with SmaI, and the digested plasmid was mixed with each pbp gene fragment separately in an ITA reaction. The reactions were transformed into *E. coli* to generate plasmid pDP563 (*P_tac_-GST-pbpD amp*) and pDP564 (*P_tac_-GST-pbpH amp*). Plasmid isolates were confirmed to have the predicted insertion by digestion with EcoRI and BamHI.

Plasmids were transformed into *E. coli* from New England Biolabs (C2566) following the included protocol. Transformations were confirmed by diagnostic EcoR1 digestion (NEB). PBP constructs were expressed in *E. coli* followed by experimental alkaline shock. PBP constructs were extracted from *E. coli* cells by incubation with 10 mg/mL lysozyme at 37 °C for 30 min followed by lysis lysed on ice with a Hielscher vial tweeter UP200st (70% C, 95% A, 5% adjustment snap) for 10 min with 30 s intervals with 30 s cooling in between. Lysates were treated with NaOH then labeled with Bocillin-FL. Samples were prepared for SDS-PAGE analysis as described above.

### Software

Expasy ProtParam was used to predict the molecular weight and isoelectric point of the PBPs (51). NIH Protein Blast was used to compare sequence coverage and identity similarities between PBPs (52). *B. subtilis* 168 was used in Protein Blast and Uniprot comparisons as this strain is the closest relative of *B. subtilis* 3610 (53).

## Data availability

All the data produced for this work are contained within the article and the supporting information

## Supporting information

This article contains supporting information (Figures S1–7 and Tables S1–6) (58–60). (16, 17, 28–30, 54–56)

## Acknowledgements

This material is based upon work supported by the National Science Foundation under Grant No. CHE-2001611, the NSF Center for Sustainable Nanotechnology. The Center for Sustainable Nanotechnology is part of the Centers for Chemical Innovation Program. SLM gratefully acknowledges the support of the University of Minnesota’s Doctoral Dissertation Fellowship during the development of this work. We thank Rebecca Calvo for materials (ponA::tet construct). DBK was funded by NIH R35 GM131783. BioRender was used to generate the TOC image.

## Abbreviations

PBP: penicillin-binding protein
HMW: high molecular weight
LMW: low molecular weight

