## Supporting Information Document for "Penicillin-binding protein redundancy in *Bacillus subtilis* enables growth during alkaline shock"

**Table of Contents**

Figure S1 Representative gels and viability of *B. subtilis* during acid shock.
Figure S2 The time required for PBP profile changes during alkaline shock.
Figure S3 The activity of *B. subtilis* PBPH and PBP4 constructs during alkaline shock.
Figure S4 Representative SDS-PAGE gel image for titration of *B. subtilis* 3610 cells over the concentration range of basic solutions.
Figure S5 The effect of buffers and osmolarity on the PBP profile generated during alkaline shock.
Figure S6 Microscopy and viability after alkaline shock.
Figure S7 Representative microscopy images.

Table S1 Experimentally determined pH optima of PBPs in comparison to their predicted pI and homology to any *B. subtilis* PBP.
Table S2 *B. halodurans* PBP information and homology to *B. subtilis* PBPs.
Table S3 The pH of the tested solutions with various concentrations of NaOH.
Table S4 Bacterial strains used in this publication.
Table S4 The pH of LB media after overnight growth of PBP-null mutants.
Table S6 Primer pairs used in mutant generation and expression plasmid construction.

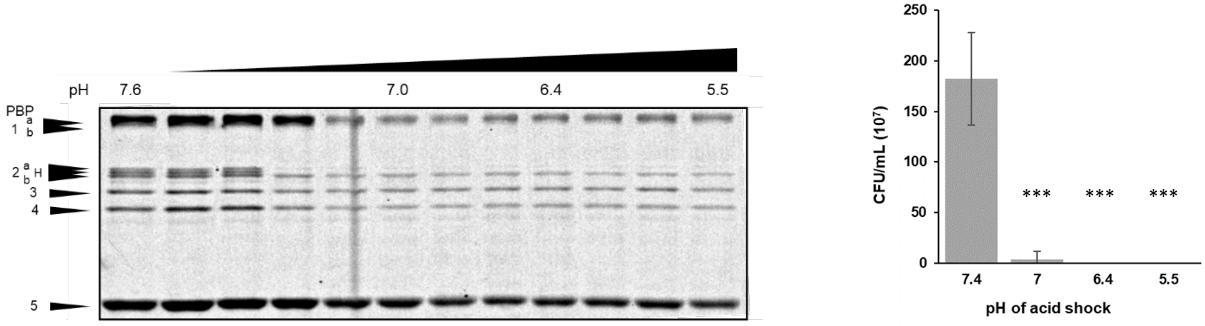


**Figure S1 Representative gels and viability of *B. subtilis* during acid shock.** (a) PBP profiles of *B. subtilis* exposed to decreasing pH in PBS for 30 min. The reduced resolution of PBPs in the SDS-PAGE gel also correlates with 99% microbial death at pH 7.0 (b) Colony forming units for bacteria incubated at noted pH for 30 min. These pH values were achieved by incubation with 0, 20, 80, and 140 mM of HCl. Error bars represent standard deviation of three biological replicates, each of three technical replicates. Significance was determined by an unpaired t-test in comparison to the control. *** $p < 0.0005$

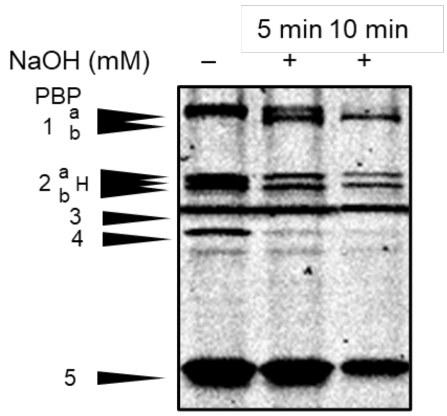


**Figure S2 The time required for PBP profile changes during alkaline shock** (pH 10.9). The "–" and "+" indicate the lack or presence of alkaline shock.

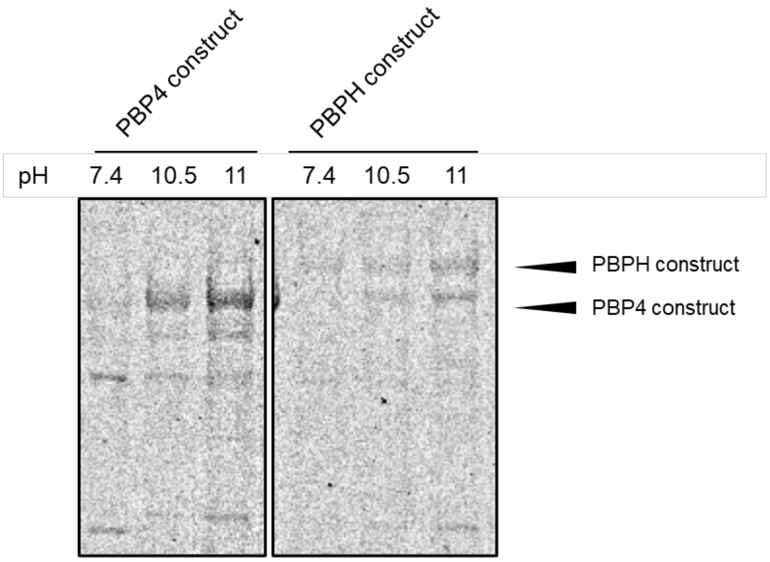


**Figure S3 The activity of *B. subtilis* PBPH and PBP4 constructs during alkaline shock** at both 4 and 8 mM NaOH, pH 10.5 and 11.1 respectively. The lysates of *E. coli*-563 containing a *B. subtilis* PBP4 construct and *E. coli*-564 containing a *B. subtilis* PBPH construct were treated.

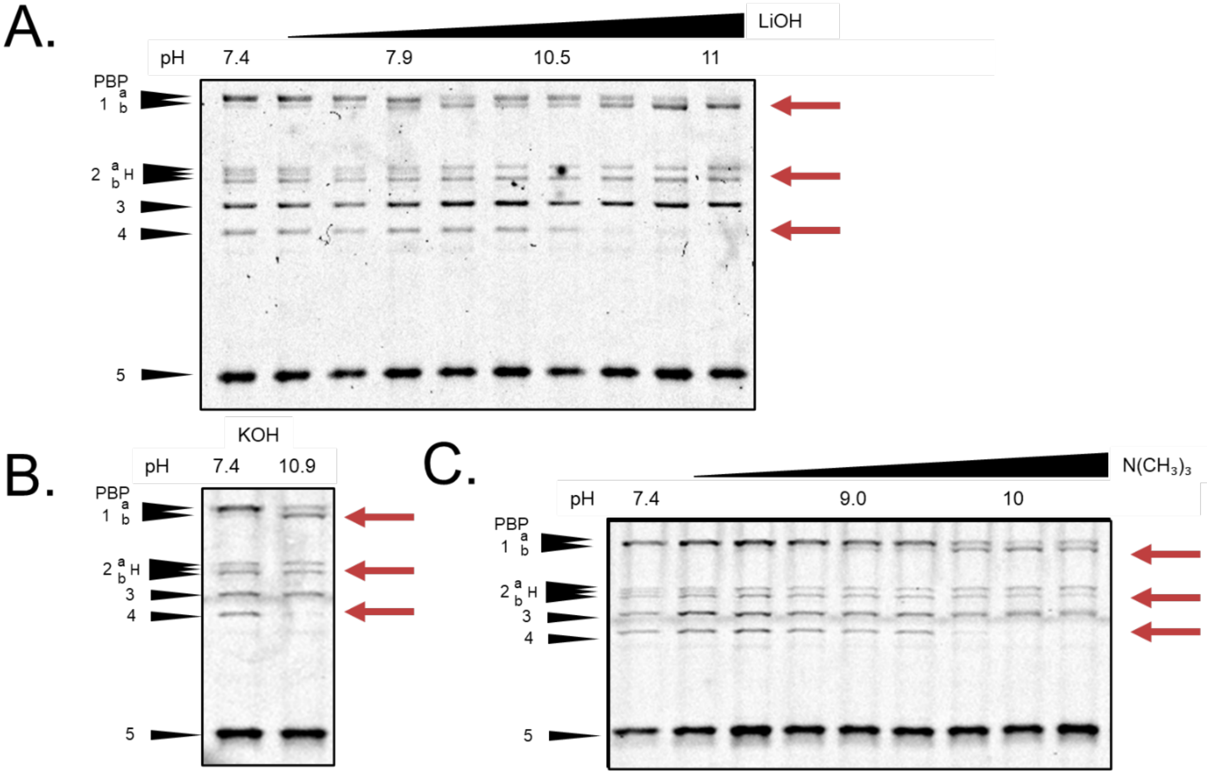


**Figure S4 Representative SDS-PAGE gel image for titration of *B. subtilis* 3610 cells over the concentration range of basic solutions**. (a) Deactivation of PBPH, PBP4, and activity shift from PBP1a to PBP1b by LiOH. (b) Deactivation of PBPH, PBP4, and activity shift from PBP1a to PBP1b by KOH. (c) Deactivation of PBPH, PBP4, and activity shift from PBP1a to PBP1b by trimethylamine. Red arrows indicate deactivation of noted PBPs.

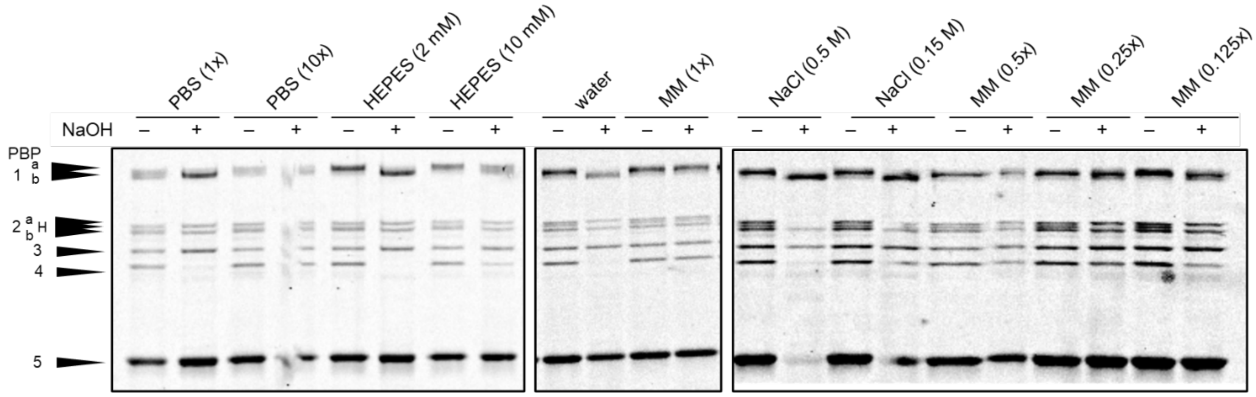


**Figure S5 The effect of buffers and osmolarity on the PBP profile generated during alkaline shock** (pH 11). The "–" and "+" indicate the lack or presence of alkaline shock. PBS: phosphate buffered saline, DPBS: Dulbeccos PBS, MM: minimal media at various dilutions. See **Table S3** for data related to the pH of each solution.

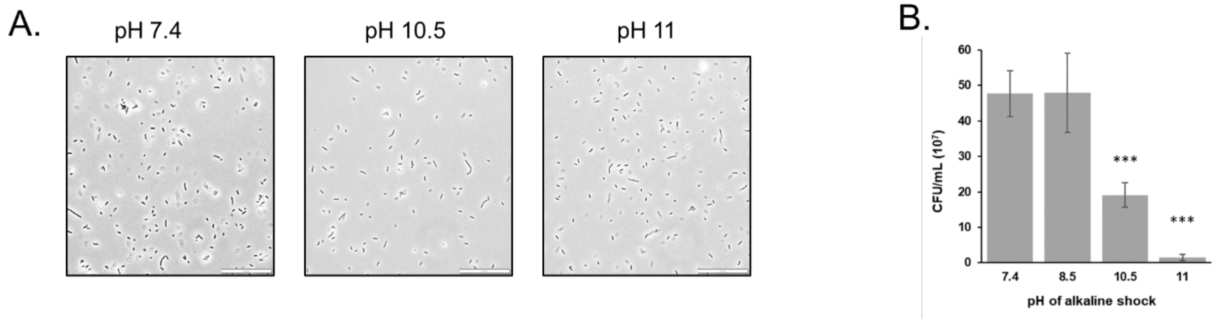


**Figure S6 Microscopy and viability after alkaline shock.** (a) Representative microscopy images for titration of *B. subtilis* 3610 cells with NaOH. Scale bar is 50 µm. (b) Colony forming units for bacteria incubated at noted pH for 30 min. These pH values were achieved by incubation with 0, 2.5, 4.0, and 8 mM of NaOH. Error bars represent standard deviation of three biological replicates, each of three technical replicates. Significance was determined by an unpaired t-test in comparison to the control. *** $p < 0.0005$.

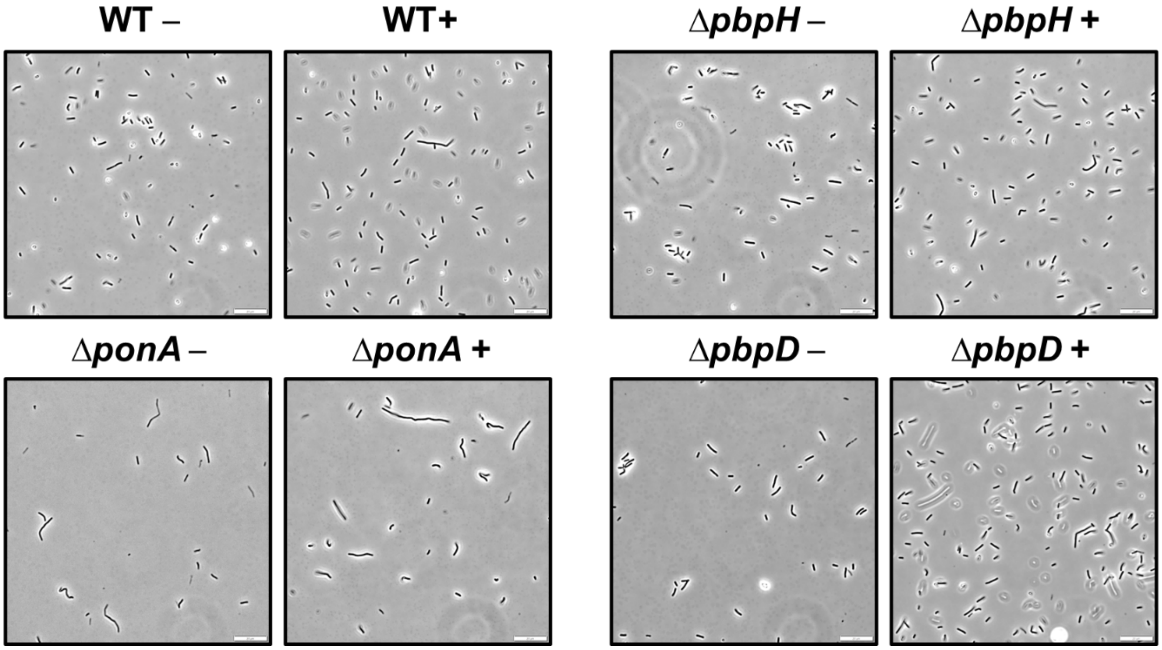


**Figure S7 Representative microscopy images** of *B. subtilis* 3610, 9744 (Δ*ponA*), DK694 (Δ*pbpH*), and 740 (Δ*pbpD)* cells with at neutral pH (**–**) and after 30 min of alkaline shock (+). The scale bar is 20 µm.

| bacteria | ID | predicted MW | predicted pI | optimal pH | lysine pKa values | homology | % coverage | % identity | reference |
|---|---|---|---|---|---|---|---|---|---|
| *B. subtilis* | PBP4a | 52891.06 | 5.53 | linear increase from pH 6 to 12 with max at pH 12 | N/A | – | - | | (1) |
| *P. aeruginosa* PAO1 | PBP5 (*pbpG*) | 34046.39 | 10.61 | 9.5 | 8.8, 10.6 | DacF (*dacF*) | 64 | 32.4 | (2) |
| | | | | | | PBP5* (*dacB*) | 71 | 28.6 | |
| | | | | | | PBP5 (*dacA*) | 72 | 26.6 | |
| | PBP2 (*pbpA*) | 72213.4 | 7.87 | 6.0-6.4 | N/A | SpoVD (*spoVD*) | 87 | 27.92 | (3) |
| | | | | | | PBP3 (*pbpC*) | 82 | 28.94 | |
| | | | | | | PBP2a (*pbpA*) | 73 | 27.5 | |
| *S. aureus* | PBP2 (*pbpB*) | 80492.48 | 8.49 | 4.5-5.0 | N/A | PBP1 (*ponA*) | 83 | 37.79 | (4) |
| | | | | | | PBP2c (*pbpF*) | 83 | 32.18 | |
| | | | | | | PBP4 (*pbpD*) | 75 | 28.03 | |
| *N. gonorrhoeae* | PBP4 (*pbpG*) | 34134.1 | 9.49 | 9 | 6.9, 10.1 | PBP5* (*dacB*) | 77 | 27.57 | (5) |
| | | | | | | DacF (*dacF*) | 83 | 27.34 | |
| | | | | | | PBP5 (*dacA*) | 73 | 27.1 | |
| *E. coli* (K12) | PBP6 (*dacC*) | 43608.91 | 7.74 | 10 | 8.4, 10.3 | DacF (*dacF*) | 85 | 34.39 | (6) |
| | | | | | | PBP5 (*dacA*) | 89 | 31.54 | |
| | | | | | | PBP5* (*dacB*) | 85 | 29.19 | |
| | PBP1a (*mrcA*) | 93636.17 | 6.15 | 7.5-8.0 | > 8 | PBP1 (*ponA*) | 78 | 39.06 | (7) |
| | | | | | | PBP2c (*pbpF*) | 59 | 36.64 | |
| | | | | | | PBP4 (*pbpD*) | 54 | 35.55 | |
| | PBP2 (*pbpA*) | 70856.61 | 8.8 | 6 | N/A | PBP3 (*pbpC*) | 85 | 26.64 | (3) |
| | | | | | | SpoVD (*spoVD*) | 94 | 25.16 | |
| | | | | | | PBP2a (*pbpA*) | 76 | 27.07 | |
| | PBP5 (*dacA*) | 44443.96 | 8.31 | 10 | 8.2, 11.1 | DacF (*dacF*) | 88 | 33.88 | (8) |
| | | | | | | PBP5* (*dacB*) | 65 | 32.34 | |
| | | | | 10 | 6.1, 9.1, 10.8 | PBP5 (*dacA*) | 89 | 30.43 | (9) |

**Table S1 Experimentally determined pH optima of PBPs in comparison to their predicted pI and homology to any *B. subtilis* PBP.**

| ***B. halodurans* protein** | **gene ID** | **MW** | **pI** | **homology** | **%coverage** | **% identity** |
|---|---|---|---|---|---|---|
| PBP1a | BH1201 | 80419.7 | 4.87 | PBP2c *(pbpF)*<br>PBP1 (*ponA)* | 94<br>80 | 52<br>35 |
| PBP2 | BH1411 | 78266.6 | 4.77 | PBP2a *(pbpA)*<br>PBPH *(pbpH)*<br>PBP2b *(pbpB)* | 95<br>91<br>64 | 44<br>40<br>24 |
| PBP1a fam | BH1702 | 98280.1 | 4.46 | PBP1 (*ponA)*<br>PBP2c *(pbpF)*<br>PBP4 *(pbpD)* | 82<br>63<br>65 | 43<br>38<br>33 |
| PBP2 | BH1894 | 78029 | 4.94 | PBP2a *(pbpA)*<br>PBPH *(pbpH)*<br>PBP2b *(pbpB)* | 95<br>91<br>64 | 41<br>37<br>25 |
| "PBP" | BH2573 | 80412 | 4.67 | PBP2b *(pbpB)*<br>SpoVD *(spoVD)* | 94<br>86 | 44<br>36 |
| "PBP" | BH3229 | 107035 | 5.07 | PBP2c *(pbpF)*<br>PBP1 (*ponA)* | 65<br>62 | 31<br>32 |
| "PBP" | BH3389 | 74827.7 | 4.16 | PBP3 *(pbpC)*<br>SpoVD *(spoVD)* | 98<br>67 | 40<br>36 |
| PBP1a | BH3812 | 77013.7 | 5 | PBP2d *(pbpG)*<br>PBP2c *(pbpF)*<br>PBP4 *(pbpD)* | 89<br>75<br>80 | 50<br>35<br>32 |

**Table S2 *B. halodurans* PBP information and homology to *B. subtilis* PBPs.**

| NaOH mM | PBS | PBS (10x) | HEPES (2 mM) | HEPES (10 mM) | MQ water | MM | MM (0.25 x) | 0.5 M NaCl | 0.15 M NaCl |
|---|---|---|---|---|---|---|---|---|---|
| **0** | 7.4 | 6.81 | 5.82 | 5.31 | 7.82 | 7.26 | 7.15 | 6.43 | 6.93 |
| **0.25** | 7.45 | 6.82 | 6.3 | 5.7 | 9.86 | 7.31 | 7.31 | 9.81 | 9.3 |
| **0.5** | 7.51 | 6.86 | 6.66 | 6.02 | – | 7.34 | 7.46 | – | – |
| **0.75** | 7.6 | 6.88 | 6.96 | 6.21 | – | 7.38 | 7.61 | – | – |
| **1** | 7.68 | 6.9 | 7.19 | 6.37 | 10.69 | 7.41 | 7.77 | 10.76 | 10.87 |
| **1.25** | 7.78 | 6.91 | 7.42 | – | – | 7.45 | 7.93 | – | – |
| **1.5** | 7.91 | 6.92 | 7.65 | 6.6 | – | 7.48 | 7.18 | – | – |
| **1.75** | 8.06 | 6.93 | 7.89 | – | – | 7.52 | 8.5 | – | – |
| **2** | 8.27 | 6.94 | 8.32 | 6.76 | 11.08 | 7.55 | 8.8 | 11.09 | 11.19 |
| **2.25** | 8.64 | 6.95 | 9.58 | – | – | 7.57 | 9.3 | – | – |
| **2.5** | 9.29 | 6.96 | | 6.89 | – | 7.61 | 9.5 | – | – |
| **2.75** | 9.65 | 6.97 | 10.29 | – | – | 7.65 | 9.65 | – | – |
| **3** | 9.97 | 6.98 | | 7 | 11.29 | 7.68 | 10.02 | 11.26 | 11.36 |
| **3.25** | 10.18 | 6.99 | 10.72 | – | – | 7.72 | 10.32 | – | – |
| **3.5** | 10.29 | 7 | | 7.11 | – | 7.77 | 10.5 | – | – |
| **3.75** | 10.39 | 7.01 | 10.94 | – | – | 7.82 | 10.61 | – | – |
| **4** | 10.49 | 7.02 | | 7.19 | 11.43 | 7.85 | 10.78 | 11.39 | 11.48 |
| **4.25** | 10.56 | 7.03 | 11.08 | – | – | 7.91 | 10.88 | – | – |
| **4.5** | 10.62 | 7.04 | | 7.29 | – | 7.96 | 10.97 | – | – |
| **4.75** | 10.68 | 7.05 | 11.18 | – | – | 8.01 | 11.04 | – | – |
| **5** | 10.73 | 7.06 | | 7.4 | 11.55 | 8.06 | 11.1 | 11.48 | 11.58 |
| **6** | 10.91 | 7.09 | 11.28 | 7.6 | 11.63 | 8.31 | 11.29 | 11.56 | 11.65 |
| **7** | 11.04 | 7.13 | 11.42 | 7.8 | 11.69 | 8.68 | 11.41 | 11.62 | 11.71 |
| **8** | 11.15 | 7.19 | 11.48 | 8.02 | 11.76 | 9.14 | 11.51 | 11.68 | 11.75 |

**Table S3 The pH of the tested solutions with various concentrations of NaOH.** The pH measured at different concentrations of NaOH. PBS: phosphate buffered saline, MM: minimal media at various dilutions.

| Species | Strain | Notes | Ref |
|---|---|---|---|
| *B. subtilis* | 3610 | WT strain | |
| | DK653 | *pbpA::kan* $comI^{Q12L}$ | this study |
| | DK654 | *dacA::kan* $comI^{Q12L}$ | (10) |
| | DK740 | *pbpD::kan* $comI^{Q12L}$ | this study |
| | DK694 | *pbpH::kan* $comI^{Q12L}$ | (10) |
| | DK695 | *pbpC::kan* $com^{IQ12L}$ | (10) |
| | DK1042 | $com^{IQ12L}$ | (11) |
| | DS9744 | *ponA::tet* | this study |
| *E. coli* (BL21) | 563 | pDP563 $P_{tac}$-GST-pbpD amp | this study |
| | 564 | pDP564 $P_{tac}$-GST-pbpH amp | this study |
| *S. pneumonia* | IU1945 | unencapsulated derivative of the D39 | (12, 13) |
| *S. aureus* | MW2 | MRSA | BAA 1707 |
| *E. coli* | K12 | | (14) |
| *S. oneidensis* | MR-1 | | BAA 1096 |

**Table S4 Bacterial strains used in this publication.**

A

| | pH | | |
|---|---|---|---|
| | **6.65** | **9** | **9.25** |
| **WT** | 6.98 | 7.67 | NG |
| **PBP 1 (Δ*ponA*)** | 6.78 | 7.67 | NG |
| **PBP2a (Δ*pbpA*)** | 6.88 | 7.96 | NG |
| **PBPH (Δ*pbpH*)** | 6.82 | 7.6 | NG |
| **PBP3 (Δ*pbpC*)** | 7.09 | 7.67 | NG |
| **PBP4 (Δ*pbpD*)** | 7.52 | 7.64 | NG |
| **PBP5 (Δ*dacA*)** | 6.75 | 7.72 | NG |

| | pH | | | |
|---|---|---|---|---|
| | **6.65** | **9** | **9.25** | **9.45** |
| **WT** | 7.87 | 7.65 | 8.05 | NG |
| **PBP 1 (Δ*ponA*)** | 7.35 | 7.65 | 7.77 | 8.28 |
| **PBP2a (Δ*pbpA*)** | 7.05 | 7.54 | 7.89 | NG |
| **PBPH (Δ*pbpH*)** | 7.27 | 7.64 | 7.72 | 8.58 |
| **PBP3 (Δ*pbpC*)** | 7.44 | 7.61 | 8.01 | NG |
| **PBP4 (Δ*pbpD*)** | 6.84 | 7.52 | 7.88 | 8.04 |
| **PBP5 (Δ*dacA*)** | 7.25 | 7.58 | 7.95 | NG |

B

| | pH | | |
|---|---|---|---|
| | **6.65** | **9** | **9.25** |
| **WT** | 7.06 | 7.5 | 8.16 |
| **PBP 1 (Δ*ponA*)** | 7.5 | 7.73 | 8.39 |
| **PBP2a (Δ*pbpA*)** | 7.59 | 8.09 | 8.31 |
| **PBPH (Δ*pbpH*)** | 7.59 | 7.9 | 8.84 |
| **PBP3 (Δ*pbpC*)** | 7.39 | 8.08 | NG |
| **PBP4 (Δ*pbpD*)** | 6.94 | 7.73 | 8.29 |
| **PBP5 (Δ*dacA*)** | 7.07 | 8.03 | NG |

| | pH | | |
|---|---|---|---|
| | **6.65** | **9** | **9.25** |
| **WT** | 7.09 | 9.07 | NG |
| **PBP 1 (Δ*ponA*)** | 7.49 | 7.75 | 8.2 |
| **PBP2a (Δ*pbpA*)** | 7.63 | 8.04 | NG |
| **PBPH (Δ*pbpH*)** | 7.98 | 8.35 | 8.56 |
| **PBP3 (Δ*pbpC*)** | 6.94 | 7.96 | NG |
| **PBP4 (Δ*pbpD*)** | 6.86 | 8.28 | 8.56 |
| **PBP5 (Δ*dacA*)** | 7.58 | 8.65 | NG |

NG = no growth

**Table S5 The pH of LB media after overnight growth of PBP-null mutants** and the ability of the PBP mutants to survive growth in basic LB media (20─30 mM NaOH generating pH 9.0─9.45 LB media), in two replicates (a) and (b). Bacteria were grown in alkaline conditions with the viable bacteria from the highest pH culture from Day 1 used to culture inoculate Day 2 cultures to assess the PBP-null mutants ability to survive in increasing pH.

| Primer | Sequence |
|---|---|
| 2973 | aggaggaattcgtgtacgtaaaaagattaaattattgct |
| 2974 | ctcctctgcagctgttataaaaaaaggatcaattttgaac |
| 3053 | aggagggatccaccatcaaatcgagcatatgaagca |
| 3054 | ctcctctgcagatctcttttccatttttatcataaatcgt |
| 3055 | aggaggaattcaaactcatcatccattgaaaaaacaaat |
| 3056 | ctcctctcgagagcttcagcaggatattaaatcaatc |
| 3250 | acgactcactatagggcgaattg |
| 3251 | ctcactaaagggaacaaaagctgg |
| 3534 | ccaagattctccgctttcag |
| 3535 | caattcgccctatagtgagtcgtcagttccacaataatccagg |
| 3536 | ccagcttttgttccctttagtgaggagctgaaatcgaagcagga |
| 3537 | gtaaactcttcggaagagaag |
| 3538 | tacagatttcaagaaaatagga |
| 3539 | caattcgccctatagtgagtcgtgaatctgttttggagcatccg |
| 3542 | gtaccagcaaagtctgagg |
| 3543 | caattcgccctatagtgagtcgtacacaaccaatgacatcagt |
| 3544 | ccagcttttgttccctttagtgagggaagcattgttgatacggta |
| 3545 | ctacaatacgaccattatgc |
| 3579 | cggaggctggtcattgtcaa |
| 3580 | caattcgccctatagtgagtcgtgagcttaaaaattaatgcagtgaa |
| 3581 | ccagcttttgttccctttagtgagacgggaacagcggaaacattt |
| 3582 | atgtcccgaagaatgacctg |
| 3583 | ccagcttttgttccctttagtgaggtatcagcctgtcgtgaaag |
| 3584 | cctgagactctaataacgaac |
| 3598 | ctaacatagctagaaaacgtc |
| 3599 | caattcgccctatagtgagtcgtggaagcgataactgtaaatgc |
| 3600 | ccagcttttgttccctttagtgaggatacaccgaccagcgttg |
| 3601 | ccatgacgacaacaaattcc |
| 7520 | gtcgtgcatctgttggatccccttccggaaaagaagtaaaacaaatg |
| 7521 | cagtcagtcacgatgaattcccttaataagccgcttgcagcgttc |
| 7522 | gtcgtgcatctgttggatcccctgaaggcgaacagcatgaagaag |
| 7523 | cagtcagtcacgatgaattcccttatttttactgtgtttttttcgagc |

**Table S6 Primer pairs used in mutant generation and expression plasmid construction.**
